# The role of reticulate evolution in biodiversity formation: the case of neotropical *Adiantum* ferns

**DOI:** 10.64898/2026.07.27.740932

**Authors:** Chi-Chuan Chen, Samuli Lehtonen, Jefferson Prado, Blake Fauskee, GoFlag Consortium, Hanna Tuomisto

## Abstract

Hybridization and introgression are thought to play key roles in the formation of biodiversity. However, detecting gene flow between species and understanding reticulation patterns remain challenging, especially in species-rich lineages with complex evolutionary histories. *Adiantum* is a large fern genus, and ecological studies in Amazonia have found that species identification is often difficult due to morphological similarity and overlapping characteristics among species. Although several hybrids have been described in tropical America, comprehensive studies investigating genetic exchanges in this genus are still lacking. We used chloroplast and nuclear phylogenomic data to examine evolutionary relationships among tropical American *Adiantum* species. By combining traditional phylogenetic analyses with advanced bioinformatic methods such as HybSeq-based target capture, reference-guided phasing (HybPhaser), and network-based analyses, we found widespread reticulate evolution involving both recent and ancient hybridization events. These were especially common among those species that have been difficult to delineate morphologically. Our findings indicate that reticulate evolution has played an important role in shaping the diversity of neotropical *Adiantum*, especially within the *tetraphyllum* lineage. Such widespread hybridization has no doubt contributed to morphological ambiguity and taxonomic challenges. Our integrated analytical approach provides a first attempt at untangling the reticulate evolutionary history in this group, and future studies with additional sampling can be expected to clarify the evolutionary processes further.

## INTRODUCTION

Ferns have been recognized as sensitive indicators of environmental and ecological variation, reflecting floristic, edaphic, and biogeographical patterns across diverse landscapes (Tuomisto & Poulsen, 1996; Ruokolainen et al., 1997; Salovaara et al., 2004; Tuomisto, 2006; Tuomisto et al., 2019). In Amazonia, some fern genera have proven particularly useful as environmental indicators, as their species distributions closely align with soil cation levels and other environmental factors (Zuquim et al., 2014; Moulatlet et al., 2019). Among these, *Adiantum* stands out as one of the most important genera, as it is both widespread and relatively species rich. Several studies have shown that *Adiantum* species compositions closely track soil and habitat characteristics, making the genus a promising ecological indicator (Tuomisto et al., 1998, 2003, 2024). However, the effectiveness of such indicator groups depends on robust species delimitation, which remains problematic in *Adiantum*. A recent study on how *Adiantum* species are distributed along an edaphic gradient in Amazonia documented the existence of 50 (morpho)species in the dataset but provided detailed information only for those 31 species that could be matched with an existing species name (Tuomisto et al. 2024).

*Adiantum* is a widely distributed genus, comprising approximately 225 species worldwide, with about 110 species occurring in tropical America (Kessler et al., 2017). The genus belongs to the early diverging fern family Pteridaceae, where it is resolved as sister to the morphologically and ecologically distinct vittarioid ferns (Lehtonen, 2011; PPG I, 2016; Pryer et al., 2016). Previous phylogenetic studies have clarified the major lineages within *Adiantum*, dividing the genus into nine well-supported monophyletic groups, with most of the Neotropical species assigned to the *peruvianum* and *tetraphyllum* lineages (Lu et al., 2012; McCarthy, 2012; Hirai et al., 2016; Huiet et al., 2018; Regalado et al., 2018). However, species delimitation within *Adiantum* remains problematic, and numerous species have been recently described, particularly from the Amazon rainforest (Prado, 2001; Lellinger & Prado, 2001; Prado, 2005; Rojas, 2007, 2008; Zimmer, 2007; Sundue et al., 2010; McCarthy & Hickey, 2011; Prado & Hirai, 2013; Boudrie et al., 2017; Prado et al., 2017).

Some *Adiantum* species have been difficult to identify both in the field and in the herbarium (e.g., Tuomisto et al. 2024). This difficulty may arise from ongoing speciation, where lineages are not yet fully differentiated, or from hybridization, where species remain largely distinct yet retain the ability to interbreed. Hybridization is especially relevant in ferns, where it is thought to occur more frequently and play a larger evolutionary role than in many other plant groups (Mallet, 2005; Sigel, 2016; Keskiniva et al. 2025). Hybridization can lead to genomic reshuffling, novel phenotypes, novel ecological traits, and enhanced competitive abilities (Baack & Rieseberg, 2007; Mallet, 2007, 2008; Chao et al., 2024; Rosser et al., 2024), but it can also increase morphological complexity and blur species boundaries.

It has been suggested that hybridization between *Adiantum* species may be relatively common, but the phenomenon has not been studied in detail (Proctor, 1985; Tryon & Stolze, 1989). At least four hybrid species have been described from tropical America, including *A.* ×*janzenianum* A.Rojas & C.Herrera, *A.* ×*moranii* J.Prado, *A.* ×*spurium* Jermy & T.G.Walker, and *A.* ×*variopinnatum* Jermy & T.G.Walker (Wagner, 1956; Walker & Jermy, 1985; Moran & Watkins, 2002; Prado, 2005; Alvarado & Martínez, 2012; Nieves & Van Ee, 2021), and others have been identified but not yet formally named (Smith, 1981; Proctor, 1985; Prado & Hirai, 2020; Castro-Aguiar et al., 2022). For some of these, at least one of the parental species is known or assumed. For instance, *A. petiolatum* Desv. has been identified a parent of *A.* ×*moranii* and *A.* ×*variopinnatum* (Walker & Jermy, 1985; Moran & Watkins, 2002; Prado, 2005), while *A. latifolium* Lam. is assumed a parent of *A.* ×*variopinnatum* and may also be involved in an undescribed hybrid (Castro-Aguiar et al., 2022).

Classical signs of hybridization are atypical morphology, failure to form sporangia, and abortive spores. Sometimes these have been observed in named species (e.g., *A. petiolatum*, *A. obliquum*, *A. lucidum*, and *A. villosum*), which might indicate hybrid origin, but it is often difficult to be sure if morphological features fall within the normal range of intraspecific variation (Castro-Aguiar et al., 2022; Lellinger, 1989; Tryon & Stolze, 1989).

The possibilities to study hybridization have improved dramatically with the development of high-throughput sequencing technologies (including target enrichment and whole-genome resequencing) and more powerful analytical methods for phylogenetic analyses. Programs such as SNaQ and HybPhaser have been successfully applied across diverse organisms to detect both historical and contemporary gene flow, clarify hybrid genomic signals, and infer parental origins (Solís-Lemus & Ané, 2016; Breinholt et al., 2021; Nauheimer et al., 2021). By integrating these approaches with genome-wide data, deeper insights can be gained into how hybridization shapes evolutionary histories and drives the diversification of complex lineages.

In the present paper, we combine chloroplast and nuclear genomic markers with advanced bioinformatic analyses to resolve taxonomic challenges in Amazonian *Adiantum*. Our objectives are 1) to assess how widespread and evolutionarily significant hybridization has been within this genus, and 2) to improve species delimitation, thereby providing a stronger genetic foundation for future ecological and biodiversity assessments.

## MATERIAL AND METHODS

### Taxon sampling

Most of the material used in this study was obtained from the collections of the University of Turku Amazon Research Team, gathered over more than three decades of extensive floristic inventories in the lowland rainforests of tropical America (Tuomisto et al., 2003, 2024; Jones et al., 2013). Voucher specimens have been deposited in at least one herbarium in the country of origin (SP and/or INPA in Brazil; QCA and QCNE in Ecuador; CAY in French Guiana; PMA in Panama; AMAZ, CUZ and/or USM in Peru) and duplicates are in the University of Turku Herbarium (TUR). Most species in the ecological dataset belong to the *tetraphyllum* and *peruvianum* lineages, so the present study focuses on the clade consisting of these two lineages.

Since most collections have served to document local species composition in community ecological studies, they have been identified to morphospecies that are thought to correspond to biological species. Most of these have been matched with a published species name by comparing with previously identified material, including type specimens. If a suitable species name was not found, a morphospecies was given a unique species number.

For the genomic analyses, multiple accessions per morphospecies were sequenced (when available) to cover their morphological variation and geographical range as well as possible.

### DNA extraction, amplification, library enrichment, and sequencing

Genomic DNA was extracted from silica-dried leaves using NucleoSpin^®^ Plant II Kit (Macherey–Nagel, Düren, Germany) following the manufacturer’s protocols. For the chloroplast dataset, the extracts were used directly for PCR amplification with four protein-coding chloroplast markers: *atpA*, *chlN*, *rbcL*, and *rpoA* (Huiet et al., 2018). The primers include ATPF412F and TRNR46F for amplifying *atpA* (Schuettpelz et al., 2006), chlN-F2 and chlN-R2 for *chlN* (Schuettpelz et al., 2016), ESRBCL1F and ESRBCL1361R for *rbcL* (Korall et al., 2006), and rpoA-F1 and rpoA-R1 for *rpoA* (Schuettpelz et al., 2016). PureTaq RTG PCR beads (Amersham Biosciences, Piscataway, New Jersey, U.S.A.) were used for PCR amplification, with different conditions for each marker. For *atpA* and *rpoA*, an initial denaturation was at 94°C for 5 minutes, followed by 35 cycles of denaturation at 94°C for 1 minute, annealing at 45°C for 1 minute, and extension at 72°C for 2 minutes, with a final extension at 72°C for 7 minutes. For *chlN*, an initial denaturation at 95°C for 5 minutes, followed by 35 cycles of denaturation (95°C, 1 minute), annealing (50°C, 1 minute), and extension (72°C, 2 minutes), with a final extension at 72°C for 7 minutes. For *rbcL*, an initial denaturation at 94°C for 3 minutes, followed by 35 cycles of denaturation (94°C, 45 seconds), annealing (52°C, 30 seconds), and extension (72°C, 90 seconds), with a final extension at 72°C for 5 minutes. The PCR products were purified and sequenced by Macrogen Inc., Seoul, South Korea/Amsterdam, the Netherlands (https://www.macrogen.com).

For target enrichment data, the concentration of extracted DNA was evaluated using the Qubit Fluorometer with dsDNA HS Assay Kit (Thermo Fisher Scientific, Oregon, USA), and DNA quality was checked by electrophoresis in 1% agarose gel. Library enrichment and high-throughput sequencing were performed at RAPiD Genomics (Florida, USA) using GoFlag 451 probes (Breinholt et al., 2021).

### Read processing, assembly, and dataset optimization

Raw sequencing reads were first processed using Trimmomatic v0.39 (Bolger et al., 2014) to remove adaptors and low-quality sequences, with the parameters: ILLUMINACLIP:NexteraPE-PE.fa:2:30:10, LEADING:20, TRAILING:20, SLIDINGWINDOW:4:20, MINLEN:30. The trimmed reads were subsequently assembled de novo using HybPiper v2.1.6 (Johnson et al., 2016), generating contiguous sequences. Two target reference files originating from the GoFlag pipeline (Breinholt et al., 2021) were used for the nuclear target enrichment dataset: an “exon-only” file containing 408 loci and an “exon plus flanking regions” file containing 410 loci.

Using the exon-only target file, 158 samples recovered 347−408 loci each. A total of 47 putative paralog loci were removed, with the mean SNP proportion across all samples being 0.02024. In addition, an average of 9 putative paralogs were removed per sample. After cleaning, the exon-only dataset included 158 samples with an average of 349 loci and a mean sequence length of 65,504 bp. For the exon + flanking regions dataset, 158 samples recovered 358−410 loci. A total of 39 putative paralog loci were removed across all samples, with a mean SNP proportion of 0.02545, and an average of 17 paralog loci removed per sample. After cleaning, this dataset comprised 158 samples with an average of 350 loci and a mean sequence length of 254,077 bp.

Additionally, a chloroplast target file was created based on the genome of *Adiantum reniforme* var. *sinense* (NC_062433) from GenBank. This file was used to retrieve chloroplast DNA sequences from the reads when available, to supplement the chloroplast DNA dataset. Summary statistics of gene recovery for each sample were generated using the command ‘hybpiper stats’, and the result was visualized using the command ‘hybpiper recovery_heatmap’ (Figure 2).

### GenBank and GoFlag data

In addition to generating new sequences for the *tetraphyllum* and *peruvianum* lineages, we used data from several other lineages to ensure a comprehensive phylogenetic analysis. The chloroplast dataset included representatives of the seven remaining *Adiantum* lineages: *digitatum*, *formosum*, *tenerum*, *pedatum*, *capillus-veneris*, *davidii*, and *philippense*, and the nuclear dataset included six of these (missing the *davidii* lineage). The additional chloroplast sequences were downloaded from GenBank, and the nuclear target enrichment data were provided by the GoFlag project team (Breinholt et al., 2021).

### Sequence alignment and phylogenetic inference for the chloroplast dataset

The concatenated chloroplast dataset included 348 samples and four genetic markers. It also integrated nine sequences assembled from GoFlag reads, with eight sequences for the *atpA* and one for the *rbcL* markers. The total sequence lengths were 1,741 sites for *atpA*, 635 for *chlN*, 1,283 for *rbcL*, and 629 for *rpoA*. In total, the dataset comprised 4,288 sites, of which 1,530 were parsimony-informative.

The DNA sequences were aligned using the MUSCLE program as implemented in Mesquite version 3.70 (Maddison & Maddison, 2021) and visually checked for ambiguously aligned regions, which were subsequently excluded. Phylogenetic trees were inferred based on maximum likelihood (ML). The dataset was partitioned by marker, and the best-fitting model was determined using ModelFinder as implemented in IQ-TREE 1.6.12 (Trifinopoulos et al., 2016). Branch supports were evaluated with 1000 replicates of ultrafast bootstrap (UFBoot; Minh et al., 2013) and Shimodaira-Hasegawa-like approximate likelihood ratio test (SH-aLRT; Guindon et al., 2010), as well as the Bayesian-like transformation of aLRT (aBayes; Anisimova et al., 2011).

### Phylogenetic inference and hybridization detection for the nuclear dataset

The results generated by HybPiper were passed to the HybPhaser pipeline, which was designed to detect the potential hybrids by assessing conflicting signals across the dataset and generating phased accessions to identify potential parental clades. The main steps of the Hybphaser pipeline included SNP assessment, Clade association, Phasing, and Merge dataset (Nauheimer et al., 2021). For the SNP assessment, we kept samples with at least 50% of loci and 30% of the target sequence length recovered, as well as loci with at least 50% of samples and 20% of the target sequence length recovered. In the clade association step, we selected reference samples for each clade that exhibited low allele divergence, low locus heterozygosity, and high locus coverage, making them unlikely to be hybrids.

Previous studies have shown that the success of HybPhaser is highly dependent on the selection of appropriate clades and reference samples to represent each clade. A sufficient number and well-distributed reference samples enable better capture of sequence variation and improve the detection of complex genomes, such as those of hybrids, whereas overly dense or sparse references may introduce noise or lead to ambiguous clade associations (Nauheimer et al., 2021). To account for this sensitivity and to identify an optimal balance, we tested several alternative clade division schemes during the clade association step. Specifically, we divided the backbone phylogeny into 13, 21, 26, 31, and 32 clades, respectively, to evaluate the accuracy and resolution of phasing outcomes for each scheme.

Following the clade association assessment, we generated phased sequences for the potential hybrids, using the two reference samples with the highest mapping rates to guide the separation of parental haplotypes. These phased sequences, together with unphased sequences representing the remaining taxa, were then merged and aligned for phylogenetic inference (Nauheimer et al., 2021).

There were two phylogenetic inferences in the HybPhaser pipeline. The first one used unphased accessions to act as a framework to decide the suitable clades that would be used for phasing, and the second phylogenetic inference used merged (unphased + phased) accessions (Nauheimer et al., 2021). For each inference, MAFFT version 7.505 was used to align the consensus sequences with default parameters (Katoh & Standley, 2013) and was then cleaned using TrimAl version 1.4.rev22 (-gt 0.5) before being passed to phylogenetic inferences (Capella-Gutiérrez et al., 2009).

We inferred coalescence-based phylogeny using IQ-TREE2 and ASTRAL-III. First, gene trees were inferred using IQ-TREE2 version 2.2.2.7 (Minh et al., 2020b) with the best-fit substitution model selected by ModelFinder and branch support assessed using 1000 ultrafast bootstrap replicates. Low-support branches with bootstrap values below 20% were collapsed to reduce phylogenetic uncertainty. The resulting gene trees were then used as input for species tree estimation with ASTRAL-III (version 5.7.8), which integrates multiple gene trees to infer a consensus species tree (Zhang et al., 2018). Support values for the species tree were evaluated using posterior probabilities, gene concordance factors (gCF), and site concordance factors (sCF) to assess the level of conflict among loci (Minh et al., 2020a).

### Assessment of reticulation and introgression

To investigate potential reticulate evolutionary relationships among the studied lineages, we applied the SNaQ (Species Networks applying Quartets) approach, implemented in the PhyloNetworks package in Julia (Solís-Lemus & Ané, 2016).

Given the computational limitations of SNaQ, we focused our analysis on the *tetraphyllum* lineage, dividing it into three clades and selecting a limited number of representative samples to form three datasets, one per clade. In addition, for suspected hybrids with potential parents from different clades, we created a fourth dataset containing samples from all three clades, but with a more limited number from each clade. Sample selection was based on the results of HybPhaser, which analyzed all samples from the *tetraphyllum* lineage. Each of the four datasets for SNaQ analysis contained no more than 30 samples, and for comparison, we also applied HybPhaser analysis to the same datasets.

Gene trees and the species tree for the SNaQ analysis were produced using the HybPhaser pipeline, with the species tree serving as the starting point. Quartet concordance factors (CFs) were estimated from gene trees to quantify genealogical discordance. Phylogenetic networks were constructed using SNaQ by iteratively increasing the number of hybridization events from 0 to 9, and 10 independent runs were used. The optimal network was primarily selected based on log-likelihood scores (Figure S1), which were compared alongside visualizations of reticulation events and inheritance probabilities to clarify evolutionary relationships.

Bioinformatic analyses of the target enrichment data for Hybphaser and SNaQ were performed using the supercomputers at CSC (Centre for Scientific Computing, Espoo, Finland). The laboratory workflow and bioinformatics pipeline used in this study are summarized in Figure S2.

## RESULTS

### Taxon Sampling

Our chloroplast and nuclear datasets represent 163 species, or approximately 70% of the currently recognized global diversity of *Adiantum*, including about 91 species that occur in tropical America. The species diversity may be underestimated, as our datasets include roughly 15–20 potential hybrids or undescribed species that require further evaluation. Detailed information on sampled accessions, represented species, and potential new species or hybrids are provided in Tables S1 and S2.

### Phylogenetic inferences for the chloroplast dataset

Our phylogenetic analysis supported the existence of nine lineages within *Adiantum*, although the *peruvianum* lineage received only weak support (SH-aLRT = 89.8% / aBayes = 0.99 / UFBoot = 100%). Among these, the *davidii* and *philippense* lineages formed a well-supported sister pair, which was in turn sister to the *capillus-veneris* lineage (SH-aLRT = 100% / aBayes = 1.00 / UFBoot = 100%). The *tenerum* and *pedatum* lineages were resolved as sisters but this relationship was only weakly supported (SH-aLRT = 57% / aBayes = 0.79 / UFBoot = 40%).

The relationships of the *digitatum* and *formosum* lineages were less clearly defined. The *formosum* lineage was basal and sister to all the other eight lineages together in our final chloroplast phylogeny (Figure 1). However, a weakly supported sister relationship between the *digitatum* and *formosum* lineages emerged both in alternative analyses of the chloroplast data (tree not shown) and in the coalescent phylogeny based on the nuclear data (Figure 2).

**Figure 1.**
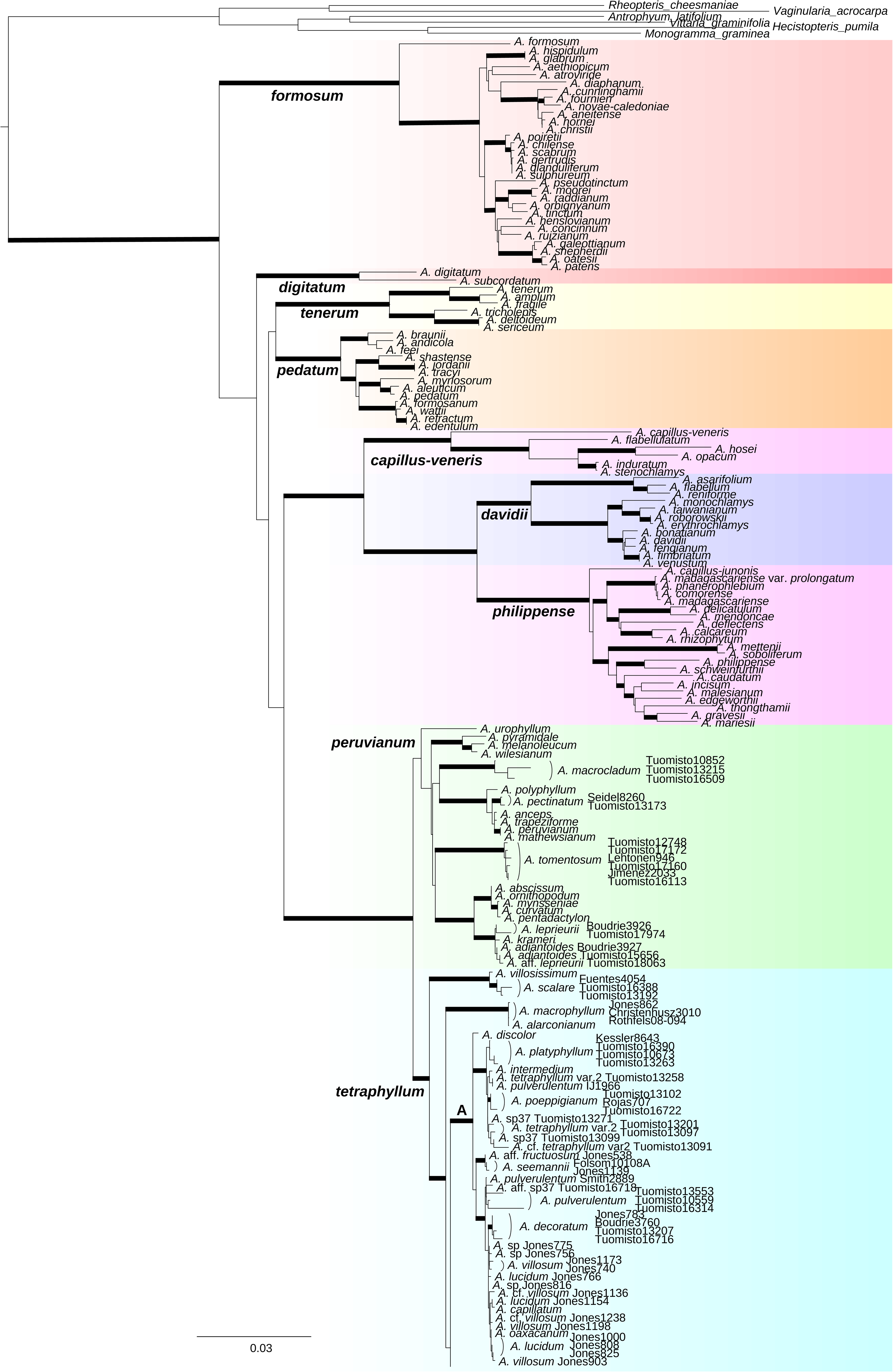
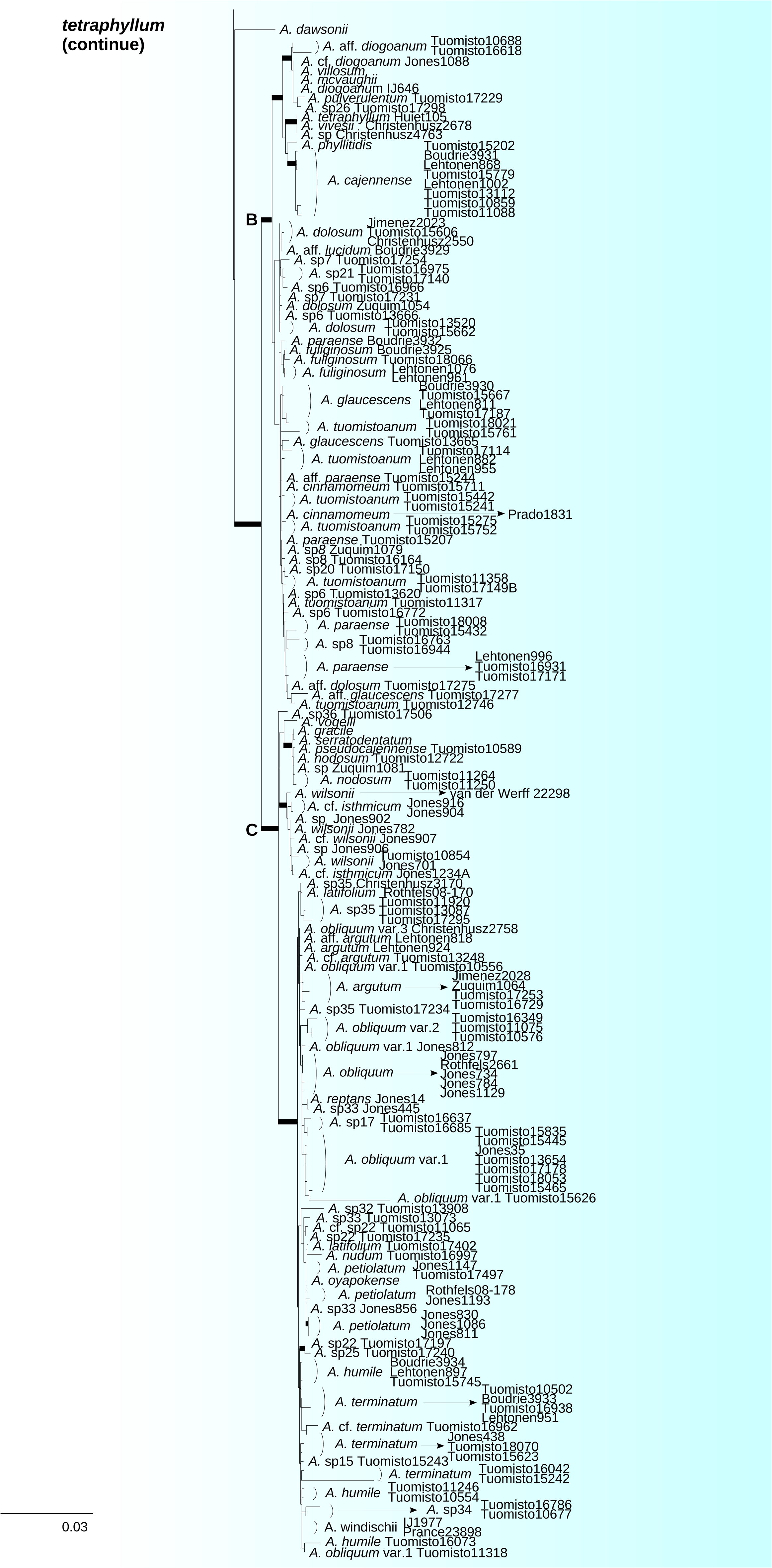
Maximum likelihood phylogeny inferred from the chloroplast dataset. The tree topology is supported by three statistical tests: SH-aLRT, aBayes, and ultrafast bootstrap (UFBoot). Bold branches indicate strong support, defined as SH-aLRT > 95%, aBayes = 1, and UFBoot = 100%.

**Figure 2.**
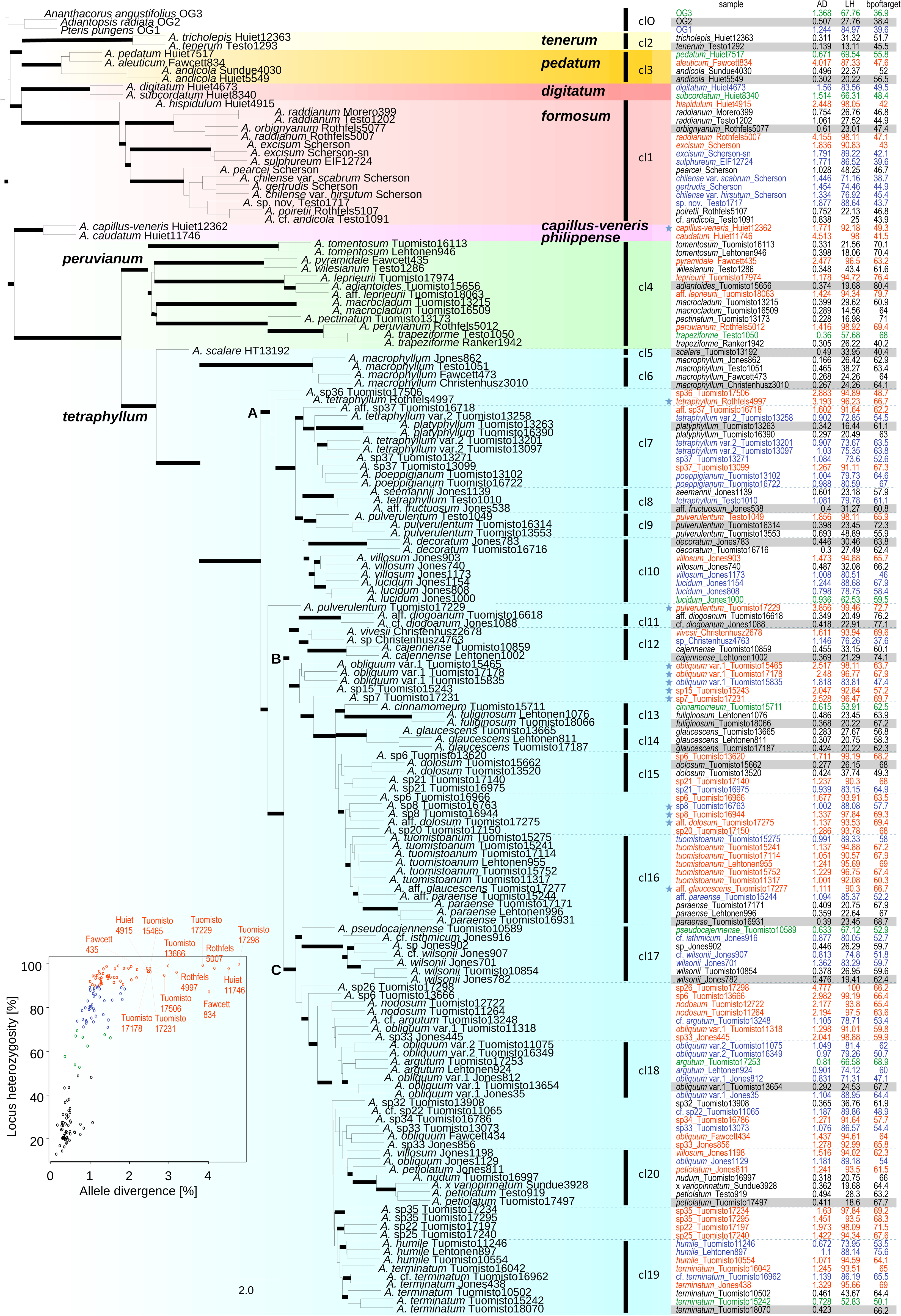
Pre-phasing phylogeny generated by the first phase of the HybPhaser pipeline, divided into 21 clades. Reference samples for each clade are shaded in grey; samples selected for phasing are marked with asterisks. Branches with posterior probabilities ≥95% are shown in bold. For each sample, locus heterozygosity (LH), allele divergence (AD), and recovered sequence length relative to the target (bpoftarget) are shown. A scatter plot summarizes LH and AD, with colors matching the table: red (LH > 90% or AD > 2%), blue (LH 70–90%), green (LH 50–70%), and black (LH ≤ 50%).

The tropical American *tetraphyllum* and *peruvianum* lineages were resolved as sisters. In our dataset, they were represented by 229 and 36 samples, respectively. Six subclades were identified within the *tetraphyllum* lineage. Three of these contain only a handful of species (the *scalare*, *macrophyllum*, and *dawsonii* subclades). The other three are more species-rich and contain most of the taxonomically problematic morphospecies: the *platyphyllum* subclade (henceforth referred to as subclade A), the *cajennense* subclade (subclade B), and the *argutum* subclade (subclade C) (Figure 1). Several named species in these subclades were not monophyletic. In particular, samples thought to represent *Adiantum lucidum, A. pulverulentum,* and *A. villosum* were split across subclades A and B; *A. dolosum*, *A. cinnamomeum*, *A. glaucescens*, *A. paraense*, and *A. tuomistoanum* were spread across subclade B; and *A. argutum*, *A. humile*, *A. latifolium*, *A. obliquum*, *A. petiolatum*, and *A. terminatum* were spread across subclade C (Figure 1, Table S1).

### Phylogenetic inferences for the nuclear dataset

The original (unphased) nuclear data recovered the same nine monophyletic lineages as the chloroplast data, but with stronger support (PP = 1), providing a well-resolved backbone topology in the coalescence-based phylogeny (Figure 2). Gene concordance factors (gCF) varied across the lineages. Among those with more than 10 samples, the *formosum* lineage had the highest gCF value (90.3, with 15 samples), while the values were lower for the two Neotropical lineages, *peruvianum* (gCF = 52.6, with 13 samples) and *tetraphyllum* (gCF = 60.8, with 117 samples). Site concordance factors (sCF) followed a similar pattern, with *formosum* obtaining the highest value (92.1), and *peruvianum* (51.8) and *tetraphyllum* (49.1) the lowest, indicating heterogeneity in the phylogenetic signal (Figure S4).

The overall topology inferred for the *tetraphyllum* lineage using the nuclear data was similar to the chloroplast topology, although it contained fewer species (e.g., the *dawsonii* subclade was not represented in the nuclear dataset) (Figure 2). Samples representing non-monophyletic species in the nuclear phylogeny were generally distributed in a similar way as in the chloroplast phylogeny, with some exceptions. The specimen of *A. pulverulentum* (Tuomisto 17229) which in the chloroplast tree was resolved to subclade B fell out of this subclade in the nuclear tree; one specimen of *A. villosum* (Jones 1198) moved from subclade A to subclade C; and three specimens of *A. obliquum* var.1 (Tuomisto 15465, 15835 and 17178) moved from subclade C to subclade B (Figures 1 & 2).

### Heterozygous signal across lineages

The exon-only dataset showed an average allele divergence (AD) of 0.76% (range: 0.05%−5.68%), whereas the dataset incorporating flanking regions had a higher average AD of 1.12% (0.14%−4.78%). Similarly, locus heterozygosity (LH) values were elevated in the flanking region dataset, with an average of 64.41% (13.11%−100%) compared to 38.98% (3.05%−94.46%) in the exon-only dataset. Since non-coding regions generally vary more, the inclusion of flanking regions increased the detection of heterozygous sites. The support values for the topology were also higher when including the flanking regions than when using the exon-only dataset. Therefore, we focus primarily on the analyses based on the exon + flanking region dataset.

Based on the exon + flanking region dataset, many samples had high locus heterozygosity (LH>90%), and twelve of these had also very high allele divergence (AD>2.4%). Although a few of the high-AD samples belonged to the *philippense* and *formosum* lineages, most were in the *tetraphyllum* lineage. Samples representing non-monophyletic species generally exhibited higher heterozygosity than samples representing monophyletic species. The highest heterozygosity was observed in *A. tuomistoanum* (LH ranging 89−97% and AD ranging 0.99−1.24%) (Figure 2), while much lower values were seen in the monophyletic *A. macrophyllum*, *A. platyphyllum*, *A. decoratum*, *A. cajennense*, *A. glaucescens*, and *A. paraense* (LH 16–38%, AD 0.17–0.47%) (Figure 2). However, some monophyletic species also exhibited moderate to high heterozygosity, such as *A. poeppigianum*, *A. nodosum*, and *A. obliquum* var. 2 (LH 79–98%, AD 0.97–2.19%) (Figure 2).

Within some species, samples differed considerably in their levels of heterozygosity. This was the case, for example, in *A. obliquum* var. 1 (LH 25–98%, AD 0.29–2.52%), *A. terminatum* (LH 20–96%, AD 0.42–1.33%) and *A. pulverulentum* (LH 23–99%, AD 0.40–3.86%). In addition, many samples that were difficult to identify based on morphological data (labelled as ‘sp.’, ‘cf.’, or ‘aff.’) exhibited high heterozygosity signals (Figure 2).

### Phylogenetic inference for the nuclear dataset based on phased sequences

Across all tested clade partitioning schemes (13, 21, 26, 31, and 32 clades), the 21-clade scheme produced the best results, with posterior probabilities, gCF, and sCF slightly higher than in the unphased tree, and the 21-clade tree was therefore used to infer the main phased phylogeny. In total, twelve samples with high heterozygosity signals (LH > 85% and/or AD > 1.5%) were successfully phased and included in the phased tree. Of these, eleven belonged to the *tetraphyllum* lineage, one from subclade A and ten from subclade B (Figure 2). Compared to the unphased phylogeny, the phased topology showed an overall increase in support values (Figure 3).

**Figure 3.**
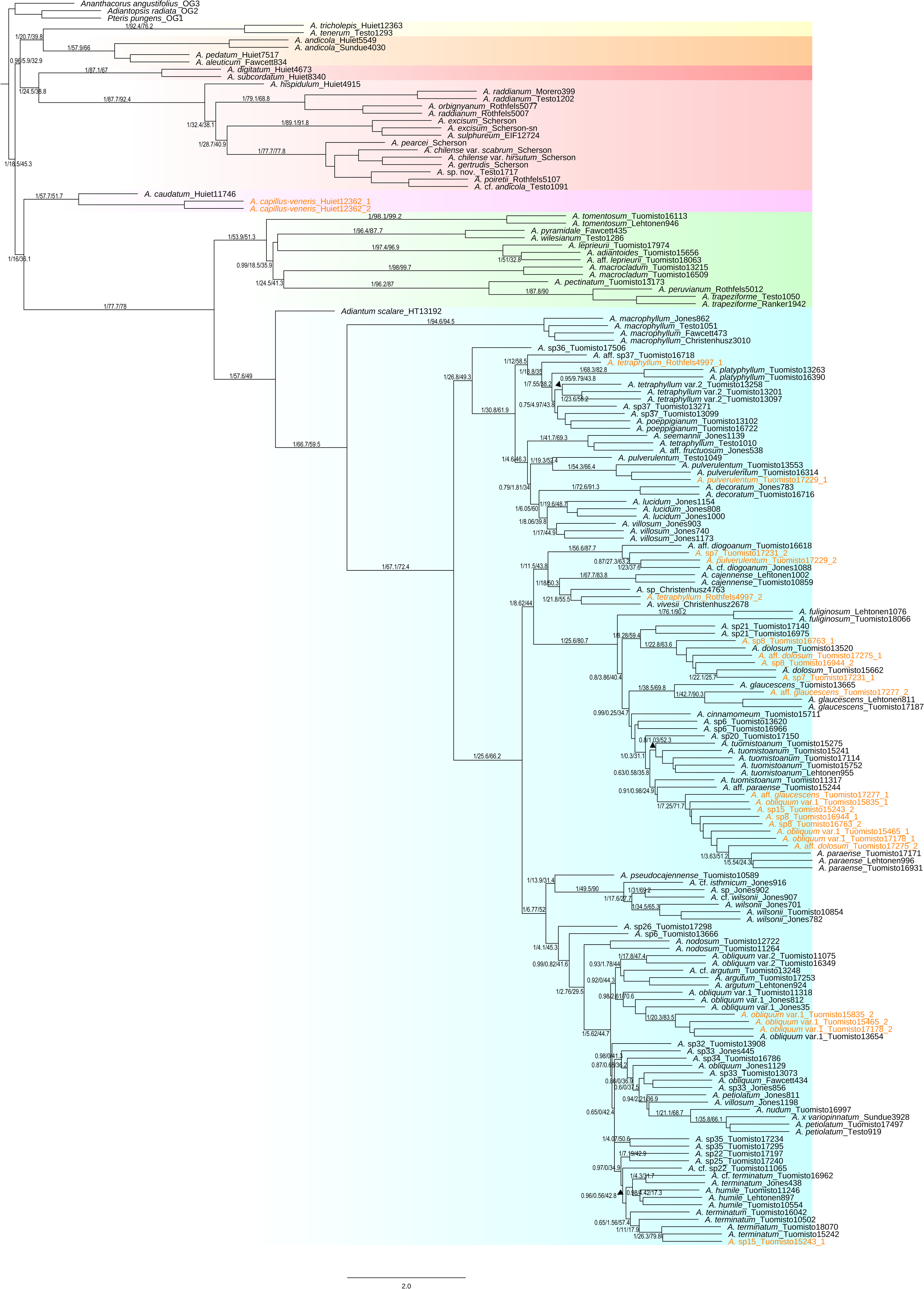
Phased phylogeny generated in the second phase of the HybPhaser pipeline. Colored tips indicate the 12 selected samples, with “_1” and “_2” denoting inferred haplotypes. Branch labels represent posterior probability / gene concordance factor (gCF) / site concordance factor (sCF).

In all eleven phased samples of the *tetraphyllum* lineage, the two haplotypes were so different that they were resolved to different phylogenetic positions. Two of the samples got split between subclades A and B (representing *A. pulverulentum* and *A. tetraphyllum*), and four samples between subclades B and C (representing *A. obliquum* var. 1 and *Adiantum* sp15) (Figure 3). Both haplotypes of the remaining five samples remained within subclade B but were resolved to separate branches within it (*Adiantum* sp7, *A.* sp8, *A.* aff. *dolosum,* and *A.* aff. *glaucescens*) (Figure 3). In general, one of the haplotypes was resolved to the clade that seemed most likely based on morphology, while the placement of the other haplotype was not always foreseen. Eight samples, representing five species, had one haplotype clustering closely with *A. paraense*, while their other haplotypes associated with four other species. One of these was *A. dolosum,* which attracted haplotypes from three different morphospecies (Figure 3).

Not all samples with high heterozygosity signals were included in the main phased analysis due to unsuccessful phasing. In some samples, allele divergence and locus heterozygosity remained high even after phasing, and separating the haplotypes did not improve the overall support values of the phased phylogeny. Therefore, they were treated as unphased data in the main phased phylogeny (Figure 3).

Although some groups showed complex reticulate relationships, our results also identified several species that formed well-supported monophyletic clades in both chloroplast and nuclear phylogenies. For example, *A. macrocladum* and *A. tomentosum* emerged as clearly delimited and well-supported species within the *peruvianum* lineage. Within the *tetraphyllum* lineage, well-supported monophyletic species included *A. scalare* and *A. macrophyllum* in basal branches; *A. platyphyllum* and *A. decoratum* in subclade A; and *A. cajennense* in subclade B (Figures 1−3). All these species showed low levels of locus heterozygosity (LH < 40%) and allele divergence (AD < 0.5%) (Figure 2).

### Network analysis and gene flow

The network analysis (SNaQ) detected several reticulation events in the *tetraphyllum* lineage, both within each of the subclades A–C and between them. In subclade A, three horizontal gene transfer events were detected. One of them was an ancient hybridization event between the ancestor of *Adiantum scalare* and the last common ancestor of the entire subclade A, and the other two events were more recent (Figure 4, A1; for the log-pseudolikelihood values, see Figure S1). The event involving *A*. aff. sp37 was paralleled by this specimen being identified as a hybrid in the Hybphaser analysis (Figure 4, A2). In contrast, the reticulation event involving *A. villosum* pointed to a sample with low heterozygosity, instead of another sample of the same species that had both high heterozygosity and was considered a potential hybrid in the Hybphaser results (Figure 4, A2).

**Figure 4.**
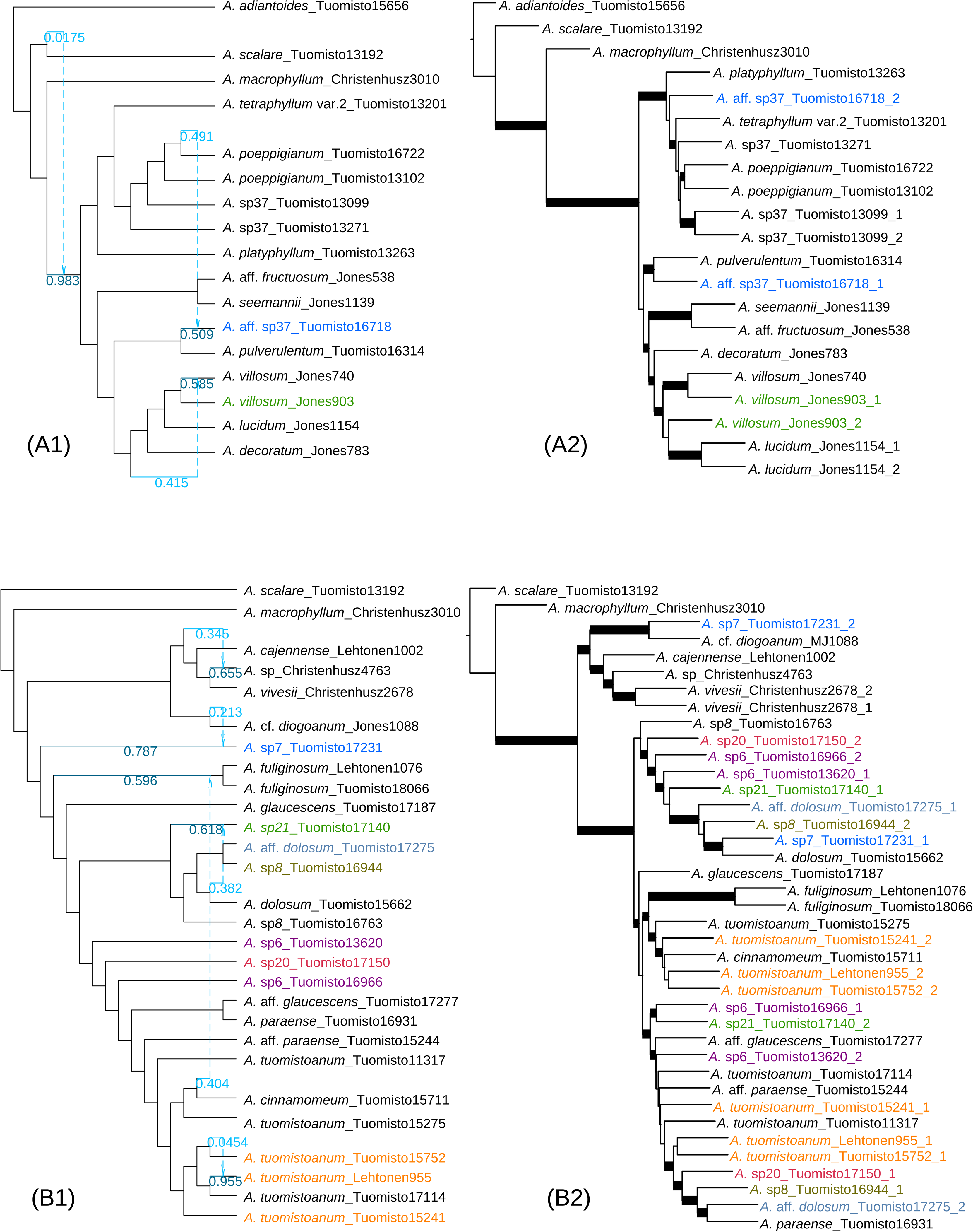
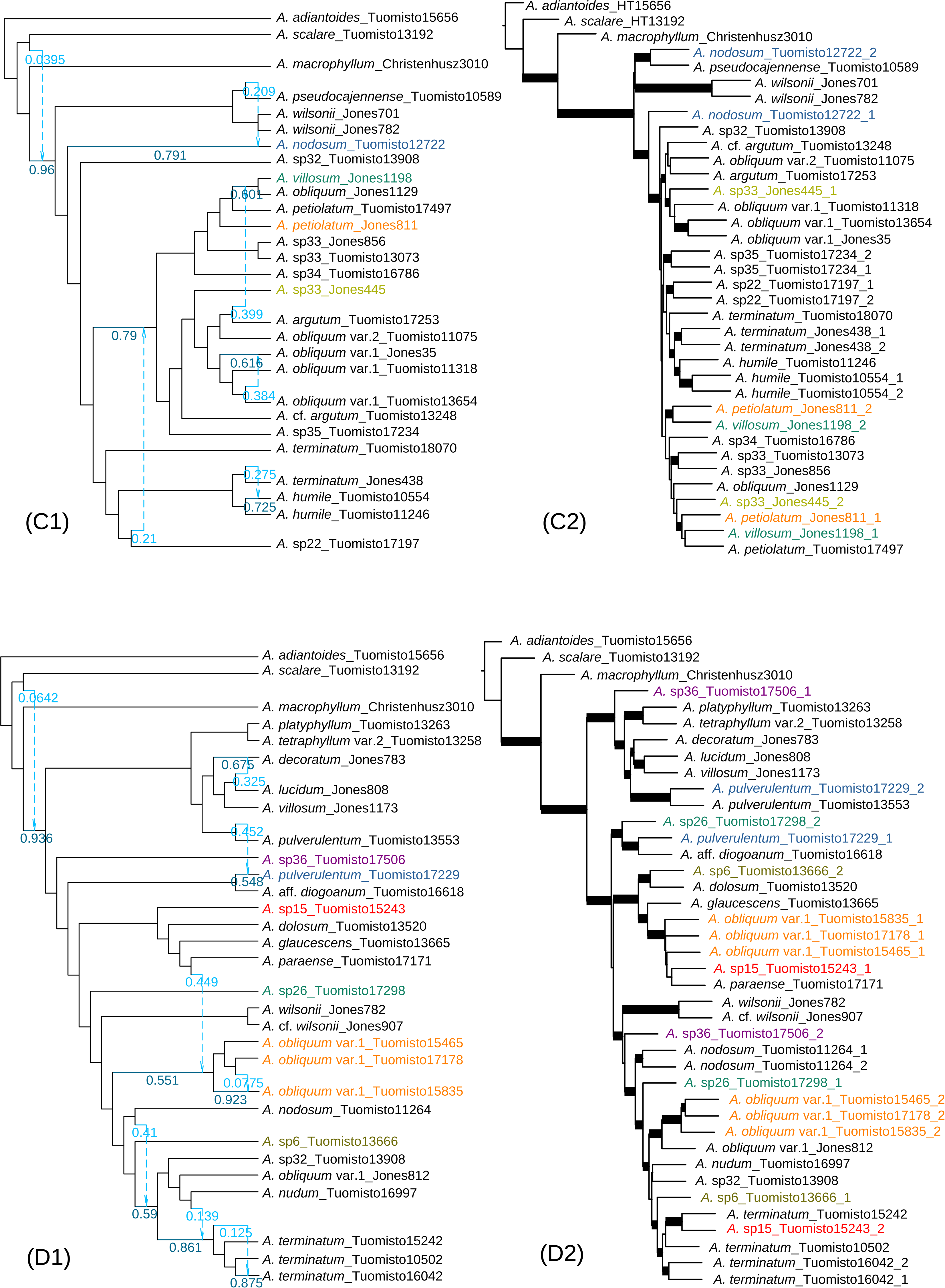
Best-supported phylogenetic networks for the *tetraphyllum* lineage inferred using SNaQ, along with HybPhaser results. The figure shows three distinct subclades (A-C) and one subclade covering the entire lineage (D). Panels A1-D1 display the SNaQ results, while panels A2-D2 show the HybPhaser results. In the SNaQ analyses, arrows indicate minor hybrid edges, with the number next to each arrow representing the inheritance probability (γ). The arrowheads point to major hybrid edges, which are labeled with 1 - γ, representing the proportion of the genome inherited vertically.

In subclade B, the best supported SNaQ result showed five reticulations (Figure 4, B1). Two of these involved samples with generally moderate to low heterozygosity, but the other three involved specimens that were identified as hybrids of *Adiantum* sp7, *A.* sp21 and *A. tuomistoanum* in the phased HybPhaser results (Figure 4, B2). However, most of the specimens whose haplotypes split into two separate branches (and therefore appear to be hybrids) in the Phased tree (Figure 4, B2) were not implicated as participants in reticulation events in the SNaQ analysis (Figure 4, B1).

Six reticulations were detected in subclade C (Figure 4, C1). The oldest <u>of these</u> involved gene transfer from the ancestor of *Adiantum scalare* to the last common ancestor of the entire subclade C. Another deep event involved gene transfer from the ancestor of *A.* sp22 to the most recent common ancestor of the *A. villosum* – *A. argutum* clade (Figure 4, C1). The more recent events appear to have transferred genes from *A. pseudocajennense* to *A. nodosum* and from *A. argutum* to the shared ancestor of *A. villosum* and *A. obliquum*, consistent with the hybrid nature of *A. nodosum* and *A. villosum* detected by HybPhaser (Figure 4, C2). Most species within subclade C showed high heterozygosity values (Figure 2), which is consistent with the deep reticulation events inferred for this group.

For the combined subclade ABC, the SNaQ network analysis inferred eight reticulation events, with the deepest one again involving *A. scalare* (Figure 4, D1). Reticulation signals were especially prominent in *Adiantum terminatum* and *A. obliquum* var. 1, both of which showed evidence of hybrid ancestry from multiple parental sources. Three samples of *A. obliquum* var. 1 also exhibited hybrid patterns in the HybPhaser analysis, but the *A. terminatum* samples did not (Figure 4, D2). With the other recent reticulations, the comparison with the HybPhaser results were mixed: in the case of *A. pulverulentum,* both a reticulation event and separation of haplotypes were inferred, but in the other cases there was indication for only one of these (Figure 4, D1–D2).

## DISCUSSION

### Hybridization and Reticulate Evolution in neotropical *Adiantum*

Our results reveal widespread non-monophyly among many morphologically defined species in the chloroplast phylogeny, while HybPhaser analyses of the nuclear dataset indicate that these species generally exhibit moderate to high heterozygosity. Reticulation analyses further suggest that hybridization and recurrent gene flow are common in Neotropical *Adiantum* and have played an important role in shaping their evolutionary history.

Three major subclades within the *tetraphyllum* lineage represent the primary regions where gene flow occurs frequently, and most described Neotropical hybrids are associated with these subclades, including *Adiantum* ×*spurium*, *A.* ×*moranii*, and *A.* ×*variopinnatum* (Jermy & Walker, 1985, 1987; Prado, 2005). Although the first two hybrids were not sampled in this study, their parental species were resolved to the subclades that show frequent reticulation. In particular, the parental species of *A.* ×*spurium* (*A. villosum* and *A. lucidum*) are associated within subclade A and exhibit clear hybridization signals (Figure 4, A2), which is consistent with previous studies indicating that these species hybridize frequently, especially in regions where their distributions overlap (Jermy & Walker, 1985, 1987). The other two hybrids (and their parental species), *A.* ×*moranii* (*A. humile* × *A. petiolatum*) and *A.* ×*variopinnatum* (*A. latifolium* × *A. petiolatum*), were primarily resolved close to each other in subclade C (Figures 1 & 2; Jermy & Walker, 1985; Prado, 2005). *Adiantum* ×*variopinnatum* occupies a basally diverging position within the clade containing one of its putative parents (Figures 2 & 3), consistent with patterns reported for hybrid taxa (Nauheimer et al., 2021).

The phylogenetic clustering of parental taxa may reflect their relatively recent divergence and high genomic similarity, which can lead to weaker reproductive barriers and a higher likelihood of successful hybridization. These patterns have been widely documented in both plants and animals, where hybridization commonly occurs among closely related lineages (e.g., Abbott et al., 2013; Mallet, 2007). However, it is also possible that the observed phylogenetic similarity does not reflect true shared ancestry, but rather the effects of historical or ongoing gene flow. Such gene flow can transfer genetic material among lineages and obscure their evolutionary relationships, leading to discordance among gene trees, particularly when occurring repeatedly over evolutionary time (e.g., Mallet, 2007; Solís-Lemus et al., 2016). Several species in our analyses exhibit such discordant patterns. For example, *Adiantum villosum* is represented by samples distributed across subclades A and C, with conflicting placements between the chloroplast and nuclear phylogenies. This pattern is commonly interpreted as the result of past hybridization and introgression, potentially leading to organelle capture (e.g., Soltis & Kuzoff, 1995; Van Raamsdonk et al., 1997; Neophytou et al., 2011; Lee-Yaw et al., 2019; Thureborn et al., 2024).

The chloroplast and nuclear phylogenies confirmed the existence of multiple genetical complexes within Neotropical *Adiantum*. Samples assigned to different species were sometimes intermingled within a single clade, while other individuals identified as the same species were resolved to divergent phylogenetic positions (Figures 1 & 2). In some species, for example *Adiantum obliquum* and *A. lucidum*, morphologically uniform samples were sometimes genetically distinct, suggesting the potential presence of cryptic species. Fišer et al. (2018) suggest that hidden diversity within taxa is often driven by factors including recent divergence, morphological convergence, and niche conservatism. Our reticulation analyses reveal both recent and ancient gene flow, with high genetic heterogeneity across most of the samples, indicating that species divergence may still be ongoing. Previously reported morphological convergence in leaf architecture (Huiet et al., 2018) further supports the role of multiple evolutionary processes in shaping cryptic diversity. Similar patterns have been reported in Amazonian trees, where species complexes are recognized even among well-known dominant species, suggesting that hidden diversity may be widespread (Damasco et al., 2021; Bacon et al., 2022; Misiewicz et al., 2023).

Our morphological identifications of specimens indirectly reflect the frequent gene flow occurring within *Adiantum*. Many individuals that were difficult to assign to known species due to ambiguous morphology were temporarily labelled as “spX” or annotated as “cf.” or “aff.” based on morphological similarity to known taxa, and these individuals tended to show high heterogeneity values. This indicates that ambiguous morphology often corresponds to genetic mixing events such as hybridization or introgression. However, not all traditionally recognized hybrid taxa display high genomic heterogeneity, as exemplified by *A.* ×*variopinnatum*. Our sampled individual of this putative hybrid shows relatively low levels of locus heterozygosity (LH = 19.68%) and allele divergence (AD = 0.36%), substantially below values typically observed in recent hybrids or introgressed individuals. This suggests that the hybrid origin of *A.* ×*variopinnatum* may reflect an ancient reticulation event followed by genomic stabilization (e.g., Buerkle & Rieseberg, 2008). Nonetheless, since only a single sample was included in this study, further sampling is necessary to determine whether this pattern is consistent across the taxon.

### Species Delimitation and Ecological Implications

Species delimitation is a fundamental component of systematics and provides the basis for describing and interpreting global biological diversity. However, species boundaries are not always clearly defined, particularly in groups where genetic exchange is frequent. Accumulating evidence shows that hybridization and introgression are widespread and often blur traditionally recognized species limits. Hybridization is far from rare; previous studies indicate that approximately 25% of plant species have hybridized with at least one other species, and the phenomenon is especially common in rapidly radiating lineages (Mallet, 2005, 2007). Our results provide clear evidence for this pattern. Frequent gene flow in Amazonian *Adiantum* likely plays a key role in its high species diversity and endemism, as it facilitates genetic variation and thereby enhances diversity across the lineage. For taxa characterized by extensive gene flow, it may therefore be more appropriate to view species as temporary and dynamic stages of divergence along an evolutionary continuum rather than as strictly discrete entities (Mallet, 2005).

Frequent hybridization and continuous evolutionary processes present a major challenge for taxonomic research, as they tend to blur species boundaries, as the morphological similarity of hybrids and backcrosses to their parental species can vary widely (Mallet, 2005). This was clearly seen in our data: several species that had been challenging to identify in the field due to intermediate forms turned out to be difficult to place also genetically. As a result, reliance on a limited number of morphological characters or a single genetic marker is unlikely to fully capture evolutionary history (e.g., Abbott et al., 2013; Huiet et al., 2018). In *Adiantum*, we observed extensive genetic admixture among subclades, discordance between nuclear and chloroplast signals, and allele sharing across lineages. These patterns suggest that species divergence in this group is both ongoing and influenced by pervasive gene flow. Earlier studies have suggested that hybridization and introgression may continue for millions of years after initial divergence (Mallet, 2005, 2007; Buerkle & Rieseberg, 2008). Rigid species delimitation in this group may be impossible, but for practical purposes one can use an integrative framework that combines morphologcal characteristics with genetic data and geographic distribution.

Why does Amazonian *Adiantum* experience such frequent hybridization? One possible explanation relates to the dynamic environmental history of the area, with various geological processes having created a multitude of different kinds of soil conditions at various spatial and temporal scales. In particular, the uplift of the Andes has changed the topography in important ways (e.g. by creating and later draining the large Pebas wetlands in the Miocene–Pliocene), and this, together with the dense network of sediment-carrying rivers has reshaped the landscape into a mosaic of soils with widely divergent chemical, mineralogical and drainage properties (Hoorn et al. 2010; Higgins et al. 2011; Quesada 2011; Tuomisto et al., 2014). Such highly heterogeneous environments provide abundant and diverse ecological niches, allowing species with different genetic backgrounds to survive and reproduce across habitats. This creates new opportunities for hybrids, enabling them to occupy niches not effectively used by their parental species and gradually form stable populations (Mallet, 2007; Moran et al., 2021; Zhang et al., 2025). These genetically differentiated populations may contribute to the potential of *Adiantum* as an ecological indicator. Several Amazonian *Adiantum* species have already been demonstrated to be effective ecological indicators (Tuomisto & Poulsen, 1996; Tuomisto et al., 1998, 2003; Zuquim et al., 2014; Moulatlet et al., 2019; Tuomisto et al., 2024). Building on this, clarifying the distribution and adaptive relationships of different genotypes across habitats will improve our understanding of their ecological roles and strengthen their application in ecological and conservation research.

### Advantages and Limitations of Analytical Methods

Our results indicate that the backbone topology recovered from the chloroplast dataset using conventional methods largely agrees with that inferred from the nuclear target-enrichment dataset analyzed with HybPhaser, as both identify nine major lineages within *Adiantum*. However, within lineages at both interspecific and intraspecific levels, the topologies produced by HybPhaser and SNaQ provide higher resolution and reveal patterns not captured in the chloroplast phylogeny. For example, closely related species within the *tetraphyllum* lineage often appeared as unresolved complexes in the chloroplast phylogeny, whereas the HybPhaser phylogeny resolved them more clearly. This likely reflects both the larger number of markers in the nuclear dataset and the fact that HybPhaser allowed analyzing the maternal and paternal haplotypes separately.

In analyses of reticulate evolution, HybPhaser and SNaQ are useful tools for investigating gene flow in *Adiantum*, but each has some limitations. For HybPhaser, we observed that many highly heterozygous samples could not be successfully phased, as these samples still exhibited extremely high heterozygosity values after phasing. The failure to achieve successful phasing in these cases may reflect underlying genomic complexity. For instance, if one or both parental lineages were themselves of hybrid origin, read mapping and allele sorting could be inaccurate, leading to misassignment or failure to assign heterozygous sites (Baack & Rieseberg, 2007; Sanderson et al., 2023). In other cases, persistent introgression or extensive reticulation may exceed the resolution capacity of the phasing algorithm, resulting in misassignments and increased noise (Browning & Browning, 2011; Tiley et al., 2024). Alternatively, some genomic regions may retain high heterozygosity from ancient hybridization events, further complicating the phasing process (Sanderson et al., 2023; Zhang et al., 2025). These scenarios highlight the limitations of HybPhaser in resolving complex reticulations. While the method is effective for detecting and phasing recent, well-defined hybridization events, it may struggle to fully capture patterns of historical gene flow and deep reticulation (Joyce et al., 2025).

In contrast to HybPhaser, SNaQ is suited for detecting both recent and ancient hybridization events across lineages and for exploring complex evolutionary scenarios. However, SNaQ is computationally intensive, and its runtime and resource requirements increase sharply with the number of taxa, making analyses with more than 30 species impractical (Hejase & Liu, 2016). To accommodate these constraints, we divided the *tetraphyllum* lineage into four datasets, including three corresponding to individual subclades and one spanning all subclades. The results indicate that all subclades within the *tetraphyllum* lineage experienced at least one ancestral hybridization event (Figure 4). Although ancestral hybridization in subclade B appears shallower than in other subclades, an alternative analysis with a different sample composition revealed a deeper reticulation within this subclade as well (Figure S3). As observed in subclade B, our broader SNaQ analyses across *Adiantum* further demonstrate that phylogenetic network inference might be sensitive to sample composition (Figure 4B1; Figure S3). Such sample sensitivity has been previously observed, especially in rapidly radiating clades (e.g., Karimi et al., 2020; Myers et al., 2024).

After comparing the reticulations inferred by the two approaches, we found that most samples that failed phasing in HybPhaser (see Results) exhibited signals of ancient reticulation in the SNaQ analyses, with several samples showing evidence of multiple gene flow events. For example, *A.* sp33 samples Tuomisto 13073 and Jones 445 each appear to have received two ancestral genetic inputs, while *A. terminatum* (Tuomisto 16042) experienced at least three ancestral reticulation events (Figure 4). Although the timing of these ancestral reticulations is currently unknown, alleles derived from ancient hybridization may be retained in descendant lineages and undergo repeated recombination. As a result, their genomic signals can no longer be clearly assigned to a single parental lineage (Baack & Rieseberg, 2007; Sanderson et al., 2023; Zhang et al., 2025). Moreover, when gene flow occurs repeatedly or continuously, heterozygosity may vary continuously, making it difficult to identify clear thresholds for phasing. As shown by our results, the distributions of locus heterozygosity and allele divergence values exhibit continuous patterns (Figure 2, inset plot).

Among the four subsets, subclade A showed the strongest agreement between SNaQ and HybPhaser. Its reticulate evolution was relatively simple, with one early ancestral event of minor genetic contribution (γ = 0.0175) and two recent non overlapping events with roughly equal parental input, allowing HybPhaser to clearly resolve relationships. In contrast, the other three subsets exhibited multiple reticulation events across both deep and recent timescales that affected different branches to varying degrees, thereby making species relationships difficult to resolve. In such complex scenarios, integrating information from both methods provides a more comprehensive reconstruction of evolutionary history. For example, in subclade B, HybPhaser showed that many alleles clustered with *Adiantum paraense* or *A. dolosum*, suggesting these species as key genetic contributors, while SNaQ inferred multiple gene flow events from them into different branches (Figure 4 & S3). Differences in gene flow strength were also reflected in the phylogeny, where taxa receiving stronger contributions (e.g., *A. obliquum* var.1) clustered more closely with the donor lineage (e.g., *A. paraense*), whereas those with weaker input (e.g., *A. tuomistoanum*) were recovered as sister lineages (Figure 3).

These findings highlight the limitations of relying on any single method. In studying the complex reticulate evolutionary history of *Adiantum*, integrating multiple analytical approaches is essential for accurately resolving hybridization patterns and gaining a deeper understanding of gene flow dynamics within the genus. Our results indicate that reticulate evolution is pervasive in *Adiantum*, involving multiple independent or repeated hybridization events that blur species boundaries. This evolutionary pattern not only explains the frequent difficulty in species delimitation based on morphology but also helps account for the observed topological incongruence between chloroplast and nuclear phylogenies in several species and species complexes.

## CONCLUSIONS AND PROSPECTS

Previous studies have indicated that tropical America is an important diversity centre of *Adiantum* (Huiet et al., 2018). In this study, we used multiple analytical methods to provide the first comprehensive evidence of extensive hybridization and introgression among *Adiantum* species in this region. Complicated reticulate evolution has led to high levels of morphological and genomic diversity, not only challenging traditional morphology-based classifications but also highlighting the need to redefine species boundaries from a more comprehensive genomic perspective.

Among the methods used, the chloroplast phylogeny was sufficient to resolve the major topological structure, even though it was based on only four markers. Therefore, at higher phylogenetic levels, chloroplast phylogeny might provide adequate resolution. However, for resolving species-level relationships, particularly in groups with a history of frequent gene flow, additional analyses using more genetic markers are necessary. In such cases, nuclear target enrichment sequencing offers a more powerful approach for resolving complex evolutionary relationships.

Our results indicate that only a limited number of neotropical *Adiantum* species exhibit low genetic heterogeneity and can be considered well-defined species, whereas most samples show varying degrees of gene flow (Figure 2). Some samples may represent populations that have recently undergone hybridization, typically exhibiting the highest heterogeneity and low gCF and sCF values. In contrast, samples with intermediate heterogeneity, such as *A. obliquum* var. 2, may represent undergoing speciation, and samples like Sundue 3928 likely represent genetically stabilized species of ancient hybrid origin. To accurately define species boundaries, future taxonomic studies should integrate both morphological and genomic information. While low heterogeneity can be a helpful indicator, species with clearly distinguishable morphological traits can also be good species, even if their genomes show some complexity. Morphology remains important for species identification, but many diagnostic traits might result from convergent evolution (Huiet et al., 2018). Integrating the phylogenies from this study with morphological data will improve the evaluation of which traits are truly reliable for distinguishing species.

Finally, hybridization is often closely associated with environmental heterogeneity. Variation in geographical and environmental factors may create opportunities for species to interact and exchange genes, and provide hybrids with opportunities to persist, resulting in variable patterns of hybridization across different habitats. Future research could benefit from incorporating time-calibrated analyses to determine when reticulation events occurred, alongside detailed assessments of habitat characteristics to evaluate their role in facilitating hybridization. These approaches would not only provide deeper insights into the mechanisms driving reticulate evolution in *Adiantum* but also help develop its hybrids as indicators of fine-scale ecological conditions.

## ACKNOWLEDGEMENTS

We are grateful to the numerous people who contributed to this study by collecting *Adiantum* specimens in the field, especially Mirkka Jones, Carl Rothfels, Layne Huiet, Michel Boudrie, and Weston Testo. We acknowledge financial support from the Turku University Foundation (grant #081252 to HT), the Research Council of Finland (grant #351460 to HT), and the Danish National Research Foundation (DNRF179 to HT).

## AUTHOR CONTRIBUTIONS

The study was initiated by SL and HT and further developed by CCC; HT, SL and JP carried out field work to collect DNA samples; HT and JP identified the species; CCC did the DNA extractions; BF contributed preliminary data and input during early discussions of the study; the GoFlag Consortium contributed additional target enrichment data; CCC conducted the data analyses with support from SL; CCC wrote the paper with major contributions from SL and HT; all authors approved the final version.

## DATA AVAILABILITY

All target enrichment sequencing reads generated during this project are available in the NCBI Sequence Read Archive (SRA) under BioProject PRJNA1466582. BioSample accession numbers and Sanger sequence accession numbers for all samples are provided in Table S1.

