## Supplementary captions for "The role of reticulate evolution in biodiversity formation: the case of neotropical *Adiantum* ferns"

**Supporting information**

**Table S1.** Summary of sampled specimens, including scientific names, assigned lineages, collection information, and genetic data from both chloroplast and nuclear sequencing.

**Figure S1.** Log-pseudolikelihood values (y-axis) plotted against the number of hybridization events h (x-axis) from SNaQ analyses for four subclades within the *tetraphyllum* lineage. The figure shows the iterative analyses across increasing h values and the number of inferred hybrid nodes (N) per h-level. The best-supported network for each subclade is marked with a bold-lined circle.

**Figure S2**. Laboratory workflow and bioinformatics pipeline used in this study.

**Figure S3.** SNaQ network showing reticulate evolution in subclade B of the *tetraphyllum* lineage within *Adiantum*, based on 26 samples.

**Figure S4.** Pre-phasing phylogeny from the first phase of the HybPhaser, with the same topology as Figure 2. Branch support is shown as posterior probabilities (PP), gene concordance factors (gCF), and site concordance factors (sCF).
