## Supplementary figures and images for "The role of reticulate evolution in biodiversity formation: the case of neotropical *Adiantum* ferns"

### Supplemental Figure S1

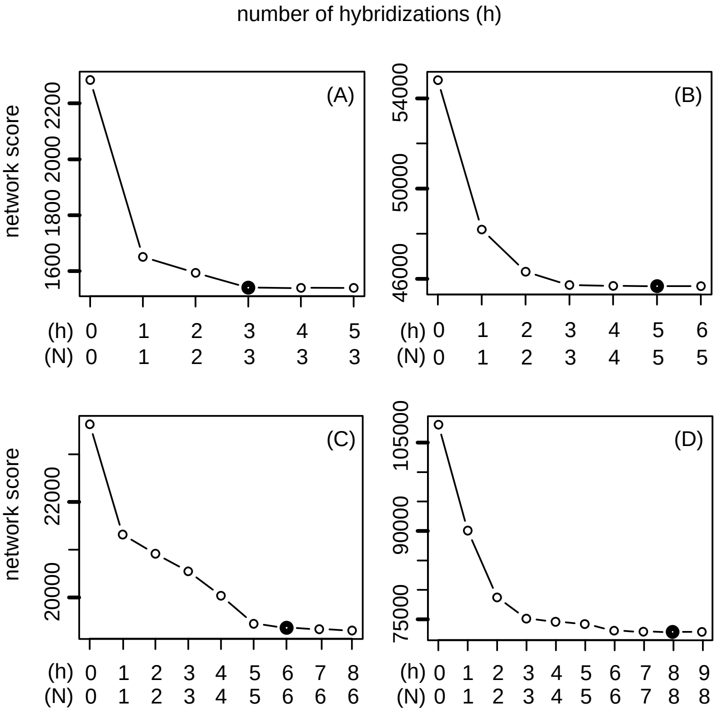

### Supplemental Figure S2

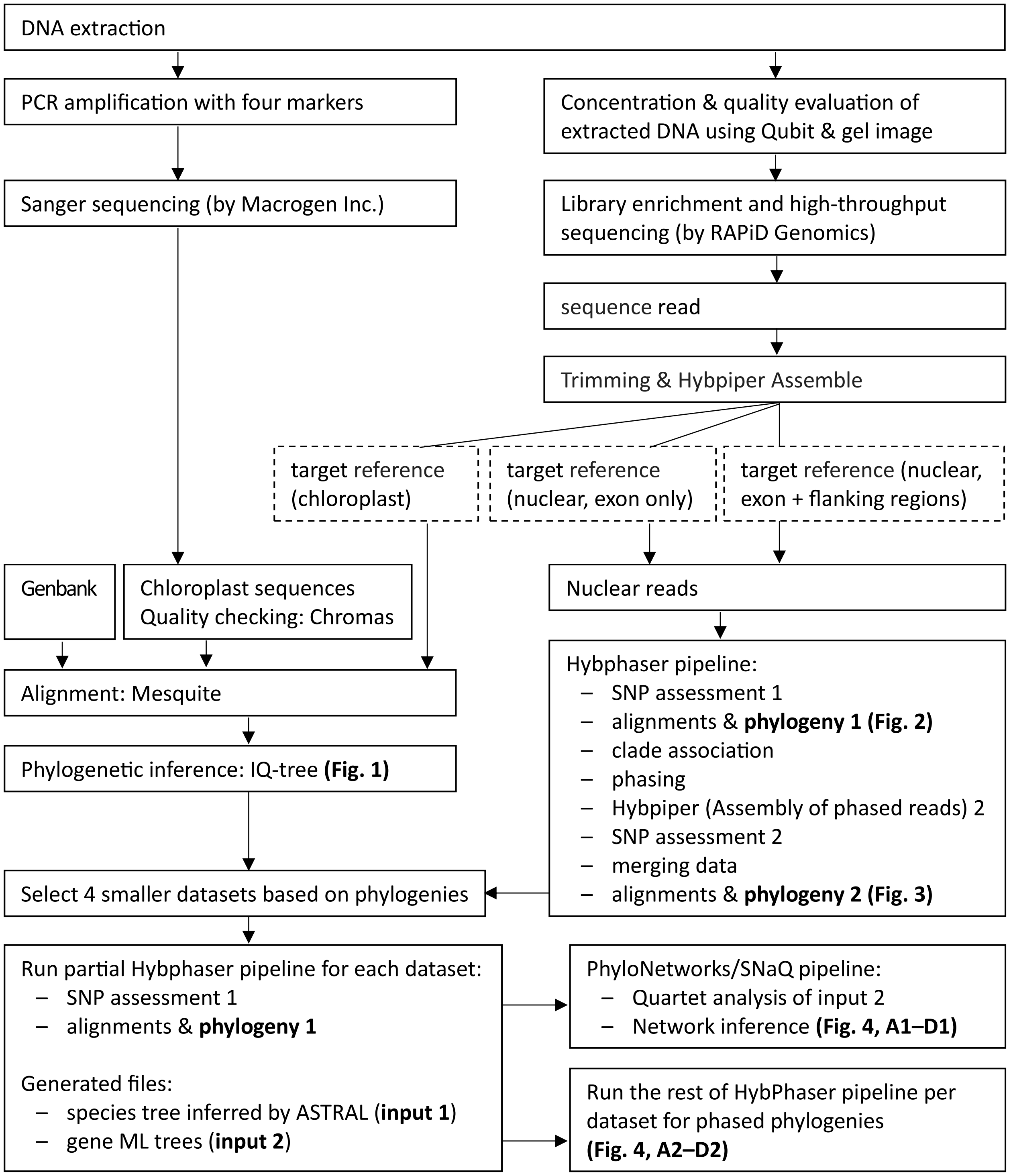


Figure S2. Laboratory workflow and bioinformatics pipeline used in this study.

### Supplemental Figure S3

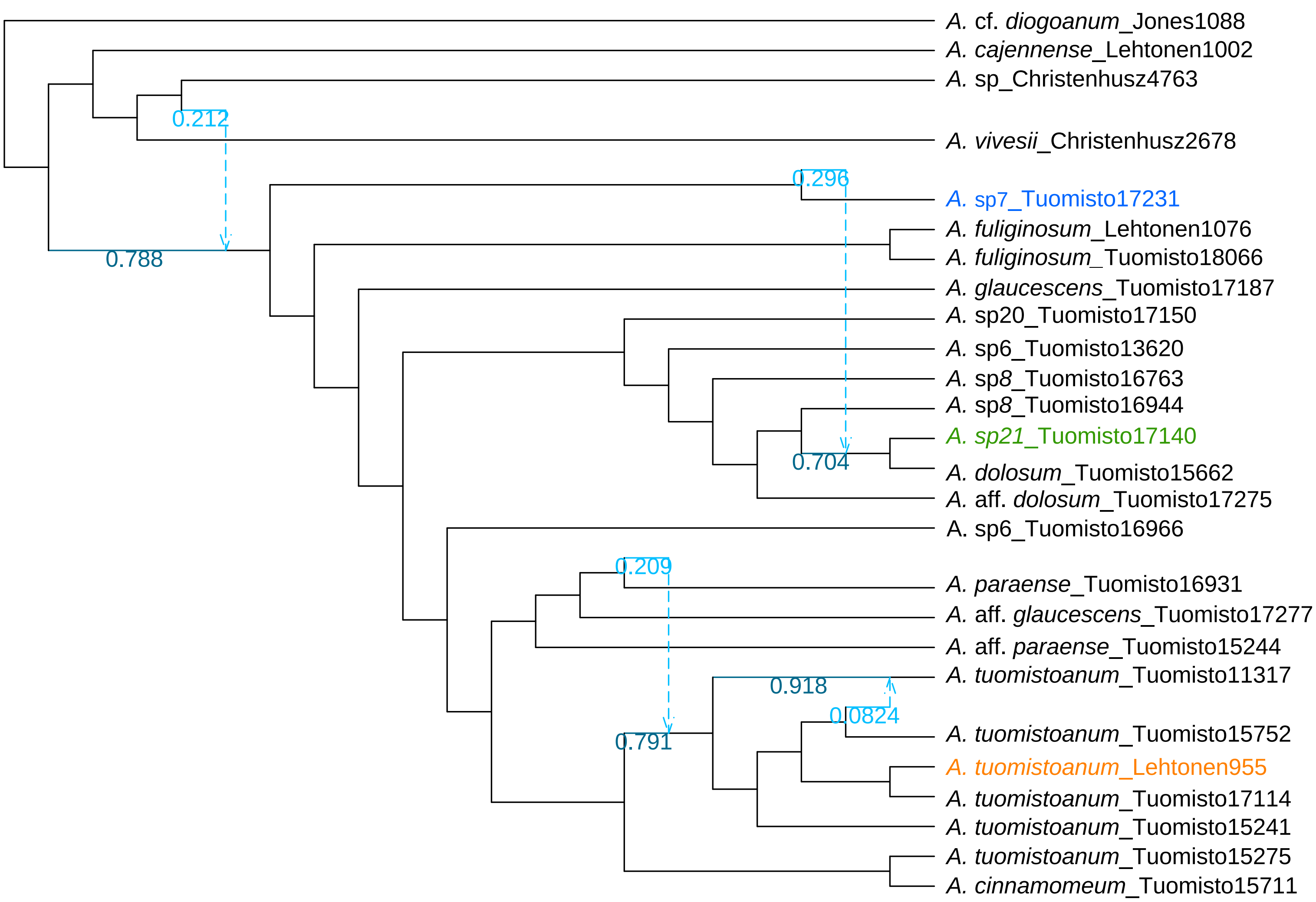

### Supplemental Figure S4

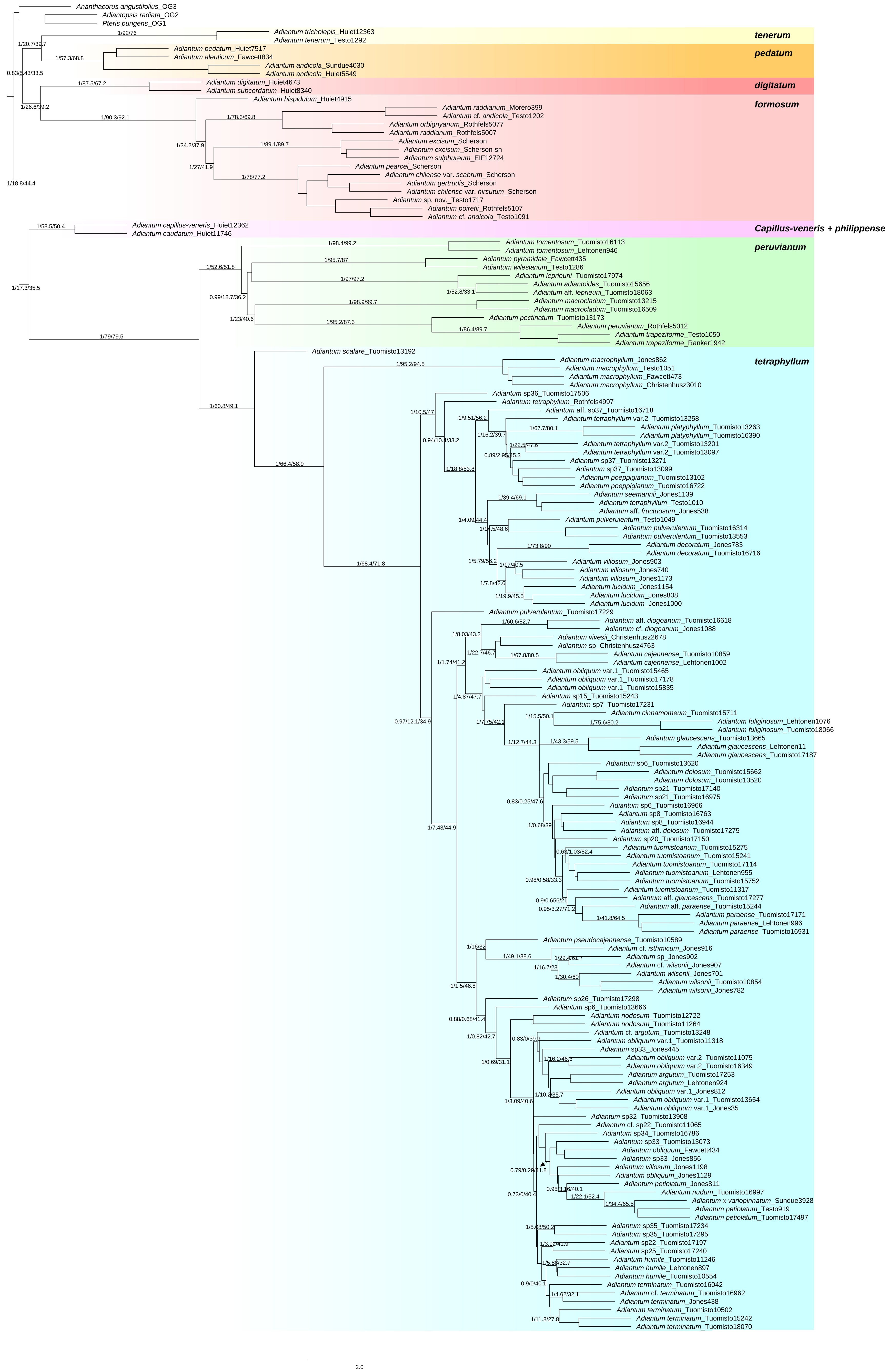
